# Brain Activity During Constraint Relaxation in the Insight Problem-Solving Process: An fNIRS Study

**DOI:** 10.1101/2022.05.09.491105

**Authors:** Reiji Ohkuma, Yuto Kurihara, Toru Takahashi, Rieko Osu

## Abstract

People solve insight problems that they encounter daily with a sudden sense of ‘aha!’ to reach a solution. Chunk decomposition, which decomposes the factors of the problem, and constraint relaxation, which manipulates filters to organize the information necessary to solve the problem, are important in insight problem-solving. Although there have been many studies on brain activity during chunk decomposition, few have examined brain activity during constraint relaxation. Particularly, no study has observed the changes over time due to constraint relaxation during insight problem-solving. This study used a slot machine task as an insight problem to measure brain activity using functional Near-Infrared Spectroscopy (fNIRS), and eye movements to estimate constraint relaxation. The results showed that the right dorsolateral prefrontal cortex (DLPFC) and right superior temporal gyrus (STG) were activated over time in participants in whom constraint relaxation was induced. The right inferior temporal gyrus (ITG) of participants who were able to solve the insight problem after constraint relaxation was also activated. The analysis of brain activity during constraint relaxation in insight problem-solving may be useful for advancing research on the ‘aha!’ phenomenon and creativity.

## 1 Introduction

People use various methods to reach solutions to the problems they encounter. One such method is called insight problem-solving, in which the solution is suddenly reached during an ‘aha!’ moment (Bowden *et al*., 2005; Weisberg, 2015). A unique feature of insight problem-solving is the solver’s experience of being in a state of limbo called impasse before reaching a solution with sudden insight. From the period of impasse, the problem is unconsciously solved through sudden inspiration. At least, consciously, there is no perceived forerunner of ‘aha’ moment.

The insight problem-solving process has long been studied in psychology and neuroscience (Weisberg, 2015; Shen *et al*., 2017, 2018). According to the representation change theory by Ohlsson (1984), the process requires chunk decomposition and constraint relaxation (Knoblich *et al*., 1999). Chunk decomposition and constraint relaxation are the two different processes in insight problem-solving. Chunk decomposition helps the solver understand the problem, while constraint relaxation is involved in the process of properly decomposing chunks. Therefore, a problem may be solved by chunk decomposition alone without constraint relaxation. Chunk decomposition is done consciously while considering the problem, while constraint relaxation is done unconsciously (Terai, Miwa and Koga, 2005).

A chunk is a component of a whole; some chunks may be easier or harder to disassemble than others. For example, in an insight problem where an incorrect equation is made using matchsticks and one matchstick is moved to the correct equation, the matchsticks that make up a number must be broken down to make up another number in chunk decomposition (‘IV->VI,’ ‘X->V’). At this point, it is relatively more difficult to decompose chunk X to V than chunk IV to VI. In contrast to Ohlsson (1984), the constraint is an implicit filter that extracts the necessary information from an infinite amount of information. In the early stages of insight problem-solving, constraints lead solvers to take the wrong approach to problem-solving. Through repeated trials, the solver gradually changes and relaxes the strength of the constraints (Knoblich *et al*., 1999). For example, in the following equation constructed using matchsticks, ‘III+III=III,’ instead of moving the matchstick that expresses the Roman numeral, it is necessary to move the one that forms the mathematical symbol. In this case, the constraint to be relaxed is ‘move the matchstick expressing the Roman numeral.’

Many studies have examined the neural mechanisms of the entire insight problem-solving process without separating chunk decomposition and constraint relaxation. A study using the compound remote associate (CRA) test, a verbal insight problem, established that the right hemisphere is more involved in the process than the left hemisphere (Bowden and Beeman, 1998; Bowden and Jung-Beeman, 2003). In CRA, one must identify a common word that is a compound of the three words presented (e.g., age / mile / sand → stone age / milestone / sandstone). In the riddles task, the right hemispheric frontal and middle temporal cortex is more involved (Zhao *et al*., 2014). The remote association test (RAT) involves the right frontal and temporal regions (Knyazev *et al*., 2021). In the RAT, one must identify a common word associated with the three words (e.g., wheel / electric / high ----chair).

In a study of a visual puzzle problem involving the decomposition of Chinese characters, the early visual cortex was inactivated and the higher visual cortex was activated (Luo, Niki and Knoblich, 2006; Wu *et al*., 2009). Another study of Chinese character puzzle problems involved functional areas for procedural memory (caudate), rewarding (substantia nigra, or SN), and visual and spatial processing (Huang, Fan and Luo, 2015). The right temporal prefrontal cortex was involved in the spatial puzzle problem (Rosen and Reiner, 2017). For problem-solving tasks that involve identifying the method used in magic tricks, regions that included the basal ganglia, the insula, parietal and temporal regions, and the frontal lobes were active (Danek and Flanagin, 2019).

There have been neuroscience studies on chunk decomposition in the insight problem-solving process; however, few of them focus on constraint relaxation. In fact, we only found one study on constraint relaxation. Huang *et al*. (2018) evaluated whether constraints were relaxed by the content of the riddle responses. For example, the constrained answer to the question ‘when the boy was slowly walking across the road and a car was speeding swiftly in his direction without any sign of slowing down, why did he not panic?’ was ‘the car is very far away from him,’ while the answer based on relaxed constraint was ‘he is walking on a pedestrian bridge.’ When the participants’ brain activity were being measured using functional magnetic resonance imaging (fMRI) as they were solving the riddles, their temporoparietal junction (TPJ) was activated. In addition, their dorsolateral prefrontal cortex (DLPFC) was active in the riddle responses that were constraint relaxed. However, Huang *et al*. (2018) did not observe the time course of the relaxation of the constraint because he analyzed whether or not the constraint was relaxed, depending on the content of the answer. Constraint relaxation changes over time after thinking. To study brain activity during constraint relaxation, it is necessary to analyze constraint relaxation before and after the activity over time.

We chose to use the slot machine task, which is a discovery task, as an insight problem that provides an analysis of constraint relaxation over time. Discovery tasks require the discovery of rules hidden in the task. This task is important for relaxing the constraints for correct chunk decomposition. By simultaneously measuring eye movement, the constraint relaxation process can be captured in detail. We, therefore, recorded eye movement and brain activity while the slot machine task was being solved.

This study clarifies the constraint relaxation process in insight problem-solving from a state of impasse to a state of ‘aha!’ where a solution is discovered. We hypothesize the following points: (1) The right DLPFC is activated in those who eventually solve the task, as they immediately relax after feeling impasse to just before having the ‘aha!’ moment; (2) Brain activity differs between those who are able to solve the task and those who cannot when they feel impasse. In the groups that did not solve the task, we considered failure to relax the constraints as the reason behind their inability to solve the task. This study will provide more insight on constraint relaxation, which will ease the ‘aha!’ experience and promote the generation of creative ideas.

## 2 Methods

### 2.1 Participants

Thirty-one university students (18 males and 13 females, aged 21.5±1.14 years) were recruited at Waseda University. Of the 31 participants, one male was excluded from the analysis because he was left-handed, although the experiment was still conducted.

The experimental procedures were approved by the Ethical Review Committee of Waseda University. All the experiments were performed in accordance with relevant guidelines and regulations. All the participants provided written informed consent prior to the experiment.

### 2.2 Procedure

In this study, the participants performed two types of tasks: a slot machine task that triggered an ‘aha!’ moment (the main task) and a slot machine task that did not (the control task).They were comfortably seated on a chair in a quiet room and instructed to look at a PC screen on a desk about 40 cm away from the chair. The procedures for the experiment provided to the participants were: ‘predict the rule for the number in the ‘?’ in the slot machine,’ ‘the correct answer will be displayed when the number is entered in the ‘?’,’ ‘identify the rule in the earliest possible trial by referencing to the correct answer,’ and ‘there is no time limit on the number of trials performed, but as soon as the number is predicted, the question must be answered.’ After the procedures were explained, a questionnaire was used to determine participants’ health status, practice tasks, and mounting and calibration of the measurement device before the measurement began.

The participants were first measured in a resting state (Rest-1) for two minutes before they performed the first slot machine task. Then, after another 2-minute resting period (Rest-2), they worked on the second slot machine task, and were again given a 2-minute resting period (Rest-3). During the rest, they looked at the cross displayed on the PC. To counterbalance, half of the participants solved the main task first and the other half solved the control task first. After the functional Near-Infrared Spectroscopy (fNIRS) measurement was completed and the device was detached, participants completed a questionnaire about the task. The questionnaire for the task asked participants about the rules found in each slot machine task and whether they experienced an ‘aha!’ moment. Each trial in each task was conducted at the participant’s pace and no time limit was set.

### 2.3 Task

This study used a slot machine task as the insight problem (Miwa and Matsushita, 2000; Terai, Miwa and Koga, 2005). The task screen consists of three slot areas with rotating single-digit numbers like a slot machine, a history window below the slot areas displaying the results of the last four attempts, and an input form above the slot areas (Figure 1-A). The slot sections are called, from left to right, slot 1, slot 2, and slot 3. Before the task, the participant is taught that the numbers appearing in slot 3 follow a specific rule.

**Figure 1.**
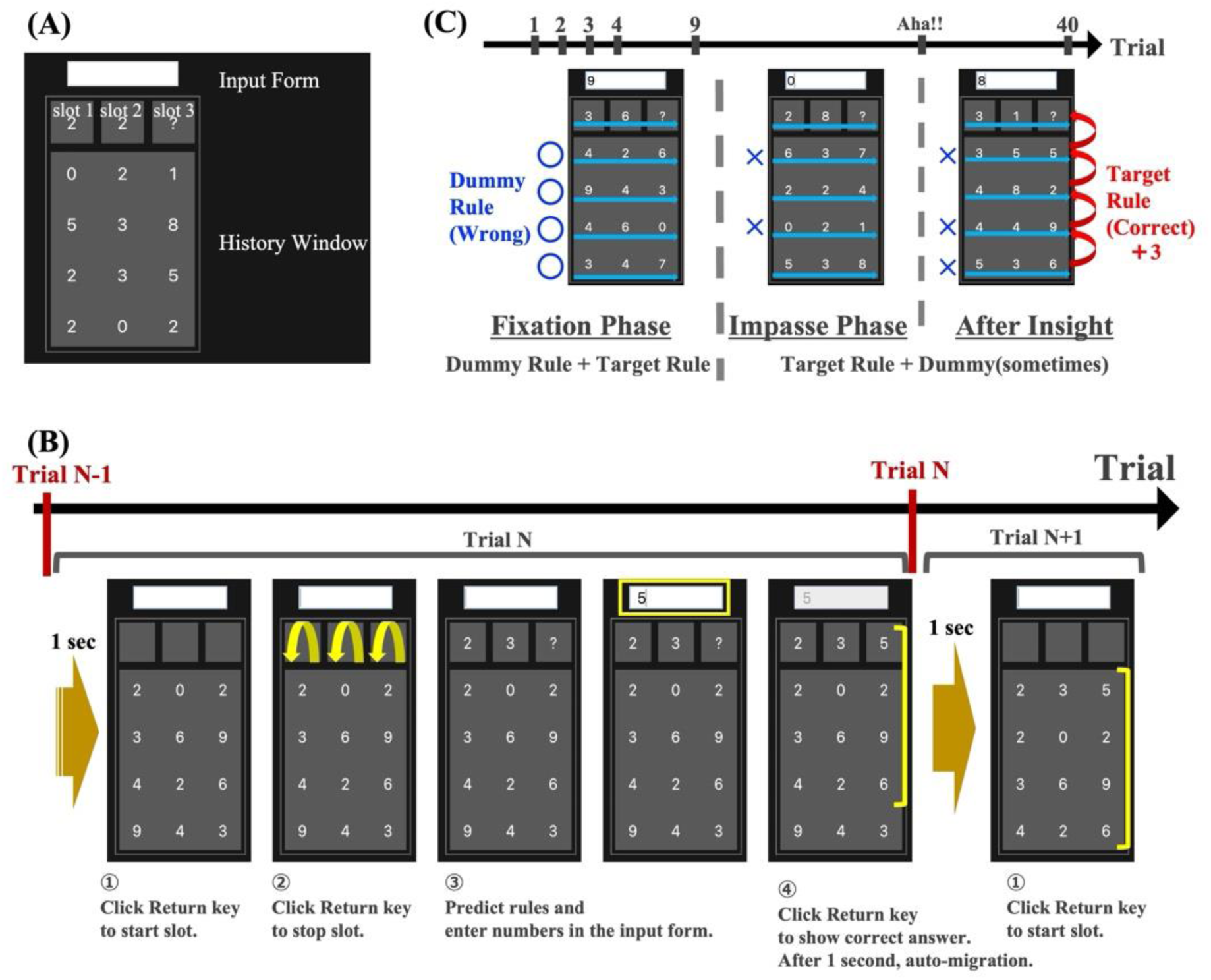
Explanation of Slot Machine Task. (A) The name of each part of the slot machine task; (B) The flow of one trial of the slot machine task; (C) Each phase of the slot machine task.

The flow of one trial is as follows. When a slot is started, the numbers in the three slots in the slot section rotate. When stopped, numbers are displayed in slots 1 and 2, while ‘?’ is displayed in slot 3. The task for the participants is to determine the rule being followed by the sequence and correctly predict the numbers that should appear in slot 3. The participant then enters the number corresponding to ‘?’ in the input form based on the rule they thought of. After answering, the correct number according to the target rule is displayed in slot 3, and the participants confirm the result. One second later, the number in the slot is automatically moved to the history window, which shows the correct numbers according to the rules, not the answered numbers. The history is only displayed for the next four trials. This process was done for 40 trials (Figure 1-B).

The rule to be determined in this task is the vertical relationship of the numbers, that is, ‘the number in slot 3 is that of the previous trial plus 3 (slot 3 vertically from 4→7→0).’ This is the target rule that is established for all trials. Apart from the target rule, a dummy rule is also established on the horizontal relationship, ‘add horizontally (2→4→6).’ The dummy rule was manipulated so that both the dummy rule and target rule were simultaneously valid from the first to the eighth trial and gradually became invalid from the ninth trial onward. In the task, the number of trials in which the dummy rule was valid was reduced from one in two to one in three, and finally to none at all. When the correct number was two added digits, the first digit was taken (Figure 1-C).

Slot machine tasks have the feature that participants have a constraint during the task and need to relax it to solve the problem. The participants tend to search for horizontal rules even after the dummy rule does not hold. As the dummy horizontal rule holds from the first to the eighth trials, the participants unconsciously have the constraint that the rule must be horizontal. To discover the target vertical rule, this constraint must be relaxed. However, even when participants receive negative feedback that rejects the hypothesis of horizontal rules in the ninth trial, they are still unable to break free from that hypothesis and often fail to find the simple target rule. In other words, this slot machine task is influenced by the dummy rule developed earlier. Unless the constraints are relaxed, the participants continue to consider incorrect hypotheses. Trials 1-8, where the dummy rule is always formed, are defined as the fixation phase, during which the participants felt stuck due to negative feedback after Trial 10. These trials are defined as the impasse phase, which is the start of the search for the target rule until the trial in which the target rule is discovered. Trials after the target rule is discovered are defined as the post-solution phase. In this study, as in previous research, we assumed that the target rule is simple enough that once it is found, a correct answer follows.

In addition to the main task which consisted of dummy and target rules, the control task was performed and only consisted of a dummy rule of lateral addition, which does not require impasse or an ‘aha!’ moment for resolution.

The success group for this task consists of participants who performed at least three trials correctly in a row after the 9th out of a total of 40 trials. The first trial of three consecutive correct responses in the success group was designated as the ‘aha!’ trial, in which the ‘aha!’ moment occurred.

Participants who had fewer than three consecutive correct answers after the ninth trial were labeled as the failure group.

### 2.4 Eye Movement Data Recording and Analyses

Eye movements were measured to estimate whether constraint relaxation occurred. In each trial, participants were judged on whether they move their eyes more horizontally based on the horizontal rule hypothesis or more vertically with the vertical rule hypothesis. Eye movement was measured using the Pupil Core (by Pupil Labs), an ophthalmic device equipped with an eye-tracking camera and subjective viewpoint camera. To wear the device, participants who needed vision correction were informed at the time of recruitment that they would have to wear contact lenses during the experiment. The sampling frequency of the eye tracking camera was 200 Hz and that of the subjective viewpoint camera was 60 Hz (720p recording). The slot machine task displayed on the PC was sized so that no more than two pieces of information overlapped in the high-resolution range defined by the retinal structure, that is, in the ‘fovea centralis,’ an area about two degrees from the center of the retina. The fonts used were small enough so they would not be difficult to see on the screen.

We only used data with a ‘config’ value of more than 0.6, which indicates reliability, to remove the blinking time when analyzing. Eye movement data for the time spent looking at the numeric keypad to enter the numbers was removed from the data. The data were split into each trial. For each one, as with Terai *et al*. (2005), we counted the number of eye movements in the ‘vertical,’ ‘horizontal,’ and ‘diagonal’ directions and calculated the ratio within that trial (Terai, Miwa and Koga, 2005). From the difference between the coordinates of the gazing point at n seconds and those of the gazing point at n+0.005 seconds (200 Hz), the difference in the horizontal direction (dx), vertical direction (dy), and the slope (dy/dx) were calculated. An absolute value of the slope greater than ✓3 was considered as ‘horizontal’ eye movement, an absolute value of 1/✓3 or less was considered ‘vertical,’ and anything in between was considered ‘diagonal’ (Figure 2).

**Figure 2.**
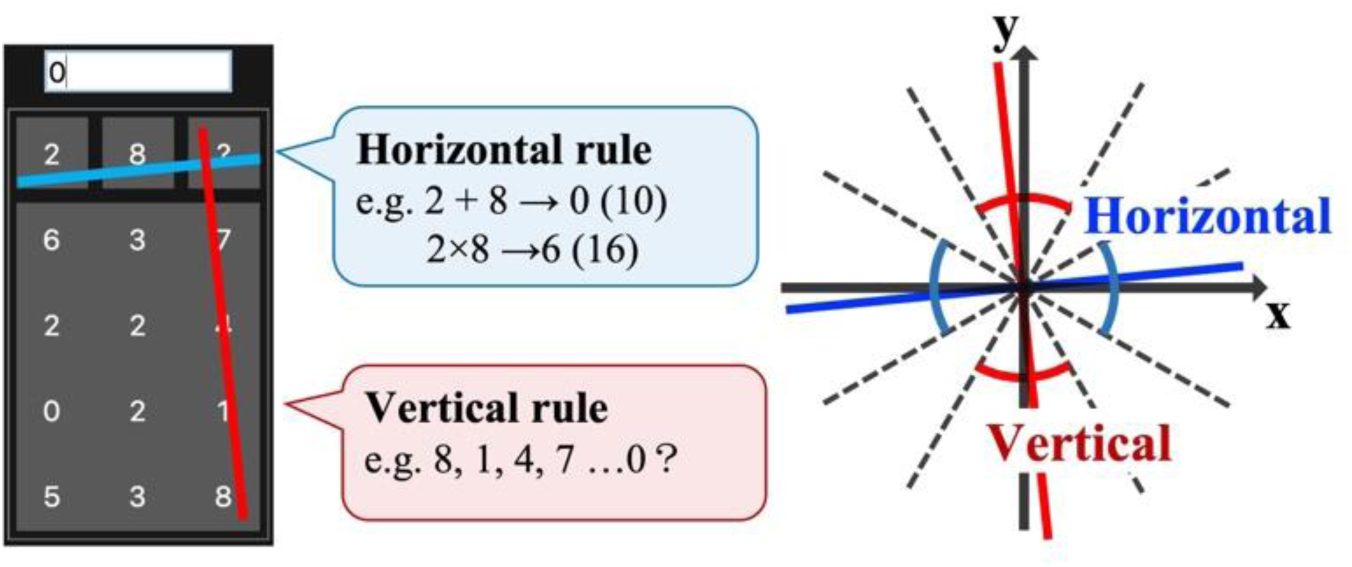
Relationship between eye movements and the hypothesized estimated rules. If the gaze point moves in the horizontal direction, the participant has a hypothesis about the horizontal direction; if the gaze point moves in the vertical direction, the participant has a hypothesis about the vertical direction. The horizontal and vertical criteria are shown on the right.

### 2.5 fNIRS Data Recording and Analyses

We used fNIRS to measure the relative concentrations of oxygenated and deoxygenated hemoglobin in cerebral blood flow to assess brain activity. NIRSports2 (by NIRx) and its measurement application Aurora were used. We used 16 source fibers, 16 detector fibers, and 16 short distance source fibers.

The short distance was used to subtract skin blood flow. The source and detector fibers were attached to the probe as shown in Figure 3. The short distance fibers were attached at the same locations as the source fibers. A total of 50 channels were measured by the probe, 34 of which were measurement channels and 16 of which were short distance pre-processing channels for analysis. The sampling frequency was 5.1 Hz. The placement of the source and detector fibers was determined using fOLD (fNIRS Optode-Location Decider) (Zimeo *et al*., 2018) based on Brodmann areas and the International 10-20 method targeting seven regions of interest (ROIs) associated with insight problem-solving processes from previous studies (e.g., Rosen and Reiner, 2017; Huang *et al*., 2018; Danek and Flanagin, 2019). The brain areas measured were the frontopolar area (BA10), orbitofrontal area (BA11), dorsolateral prefrontal cortex (BA9 and 46), inferior temporal gyrus (BA20), middle temporal gyrus (BA21), superior temporal gyrus (BA22), and angular gyrus and temporo-parietal junction (BA39) (Figure 3).

**Figure 3.**
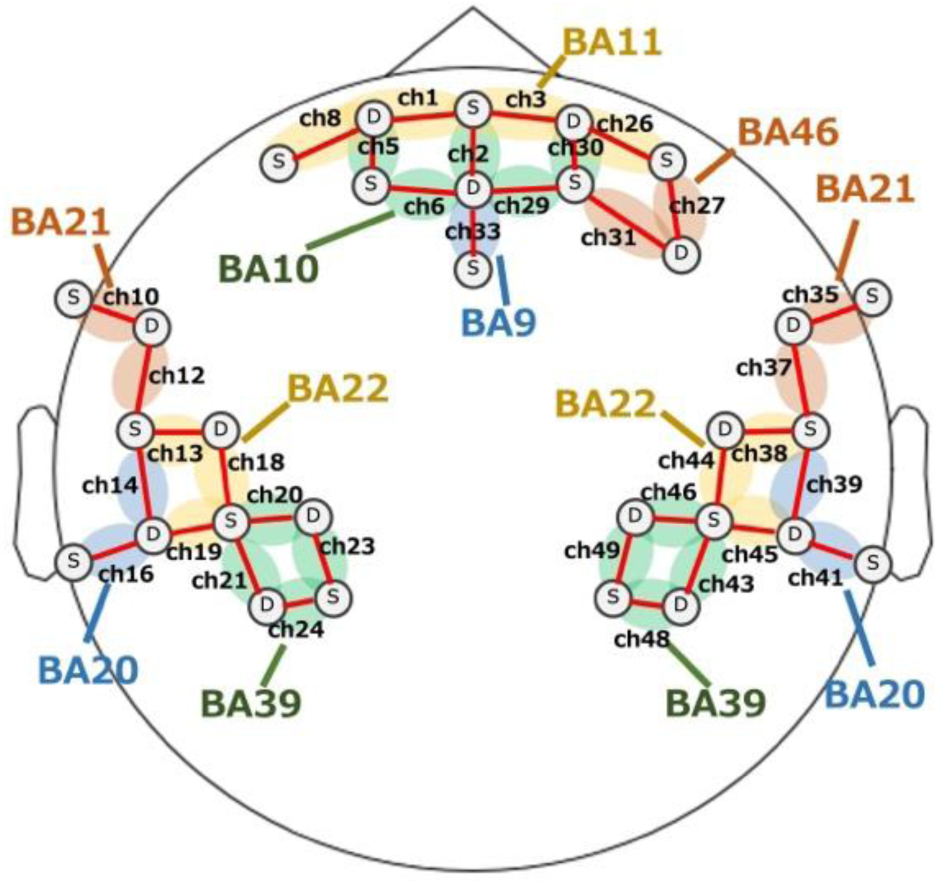
Brain regions measured by fNIRS. ‘S’ is the source and ‘D’ is the detector, measured between ‘S’ and ‘D.’ There are 34 channels with missing numbers.

We analyzed the changes in oxygenated hemoglobin concentration in the fNIRS data. First, because fNIRS data contain various kinds of noise, we performed preprocessing using open POTATo (Yamada *et al*., 2012). The preprocessing steps are as follows: 1) short distance processing (opSSR) was done to remove skin blood flow changes; 2) bandpass filtering (0.01-0.5 Hz) using an FFT (Band Filter) was performed (Tempest and Radel, 2019); and 3) the overall data for each participant was corrected with Rest-1 as baseline and all times were calculated into Z-scores for each participant (Ono, 2019). After preprocessing, the average value for 30 seconds during the task was used as the representative value for paired t-tests. For the tests, SciPy in Python was used. NIRS-MNE was used for the figure plots.

**Table 1.** ROI-channel mapping.

| **Brodman** | **Region name** | **channel** |
| --- | --- | --- |
| BA10 | Frontopolar area | ch2<br>L: ch5, 6<br>R: ch29, 30 |
| BA11 | Orbitofrontal area | L: ch1, 8<br>R: ch3, 26 |
| BA9, 46 | Dorsolateral prefrontal cortex | ch33<br>R: ch27, 31 |
| BA20 | Inferior Temporal gyrus | L: ch14, ch16<br>R: ch39, ch41 |
| BA21 | Middle Temporal gyrus | L: ch10, 12<br>R: ch35, 37 |
| BA22 | Superior Temporal gyrus | L: ch13, 18, 19<br>R: ch38, 44, 45 |
| BA39 | Angular gyrus /<br>Temporo-parietal junction | L: ch20, 21, 23, 24<br>R: ch43, 46, 48, 49 |

## 3 Results

### 3.1 Slot Machine Task

Of the 30 participants in the analysis, 15 were grouped into the success group that found the target rule and 15 were grouped into the failure group that did not find the target rule. None of the participants discovered, from the beginning, the target rule that was also correct on the ninth and tenth trials immediately after the dummy trials that were not valid. In addition, none discovered the dummy rule with incorrect answers in trials 1-8. When asked about the rule for the main task, all of the participants in the successful group selected the rule to be the target rule in the post-experiment questionnaire. In contrast, no one in the failure group answered the question correctly. The success group responded that they had an ‘aha!’ moment when they discovered the target rule. In addition, the control task was solved by all participants.

The success group found the target rule in an average of 28.9 trials (SD = 7.12). To compare whether there was a difference in the overall time spent working on the task between the success and failure groups, a t-test was conducted where no significant difference was found [t (29) = −0.2673, p = 0.7912].

For the analysis, ‘start impasse,’ ‘before aha!,’ ‘trial 29,’ and ‘control’ are defined. The participants spent more time on a trial after the ninth than the ones before. It was because they noticed that the dummy rule had failed, felt impasse, and started looking for other rules (Terai, Miwa and Koga, 2005). Therefore, the period of 30 seconds from the end of the trial (defined as trial m-1) before the longer trial (defined as trial m) (previous time ± 2 SD) is defined as ‘start impasse.’ The period immediately before the ‘aha!’ moment occurs is defined as ‘before aha!’ Specifically, ‘before aha!’ is defined as the last 30 seconds from the moment of the first trial (defined as trial n) of the three consecutive correct trials (trial n∼n+2). To study the change over time in the failure group, we defined trial 29 as the 30 seconds before the moment participants in the failure group found an answer in the 29th trial, as participants in the success group had an ‘aha!’ moment after an average of 28.9 trials. ‘Control’ is defined as the last 30 seconds during which the control task is solved without feeling impasse (Figures 4, 5).

**Figure 4.**
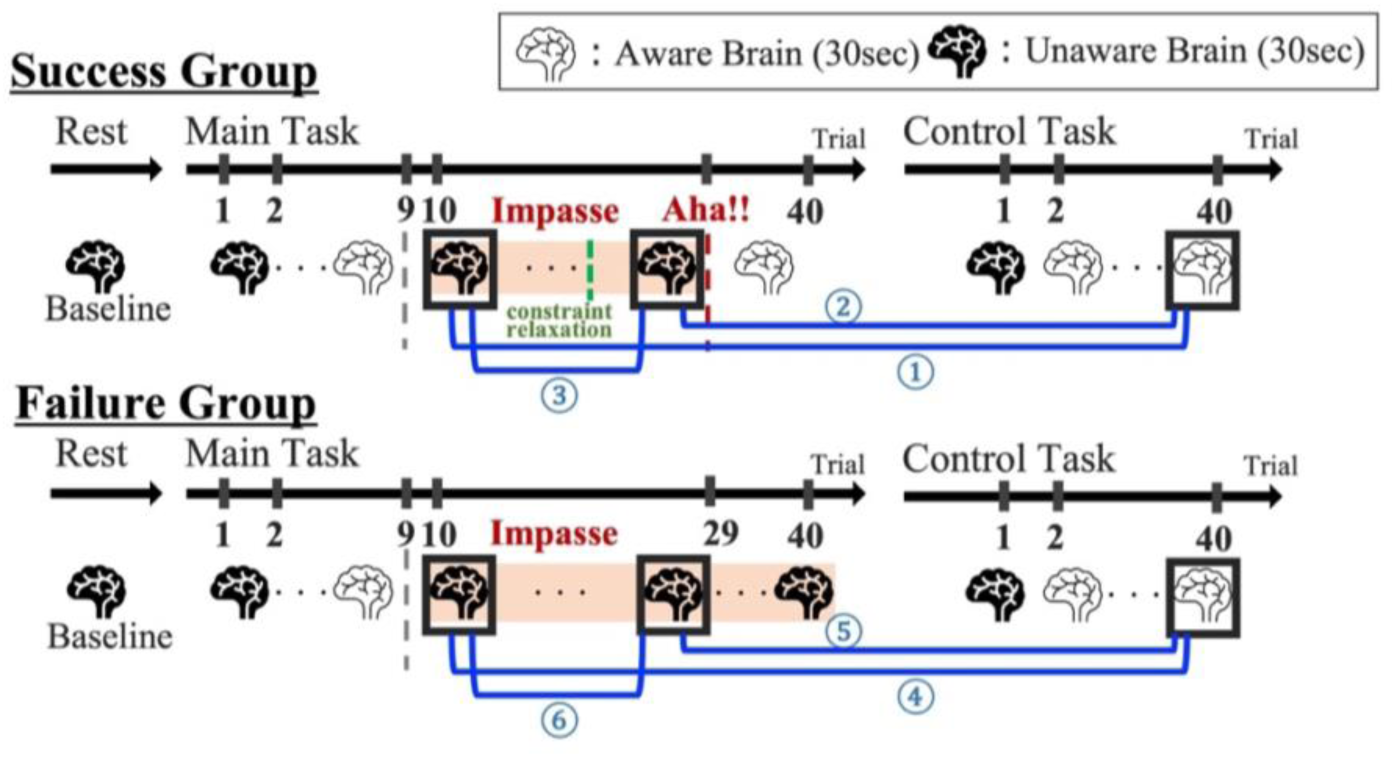
Time periods compared to clarify constraint relaxation. The parts related to constraint relaxation were cleared by comparison with the controlled task, removing the components necessary for the execution of the slot machine task and for simple calculations.

**Figure 5.**
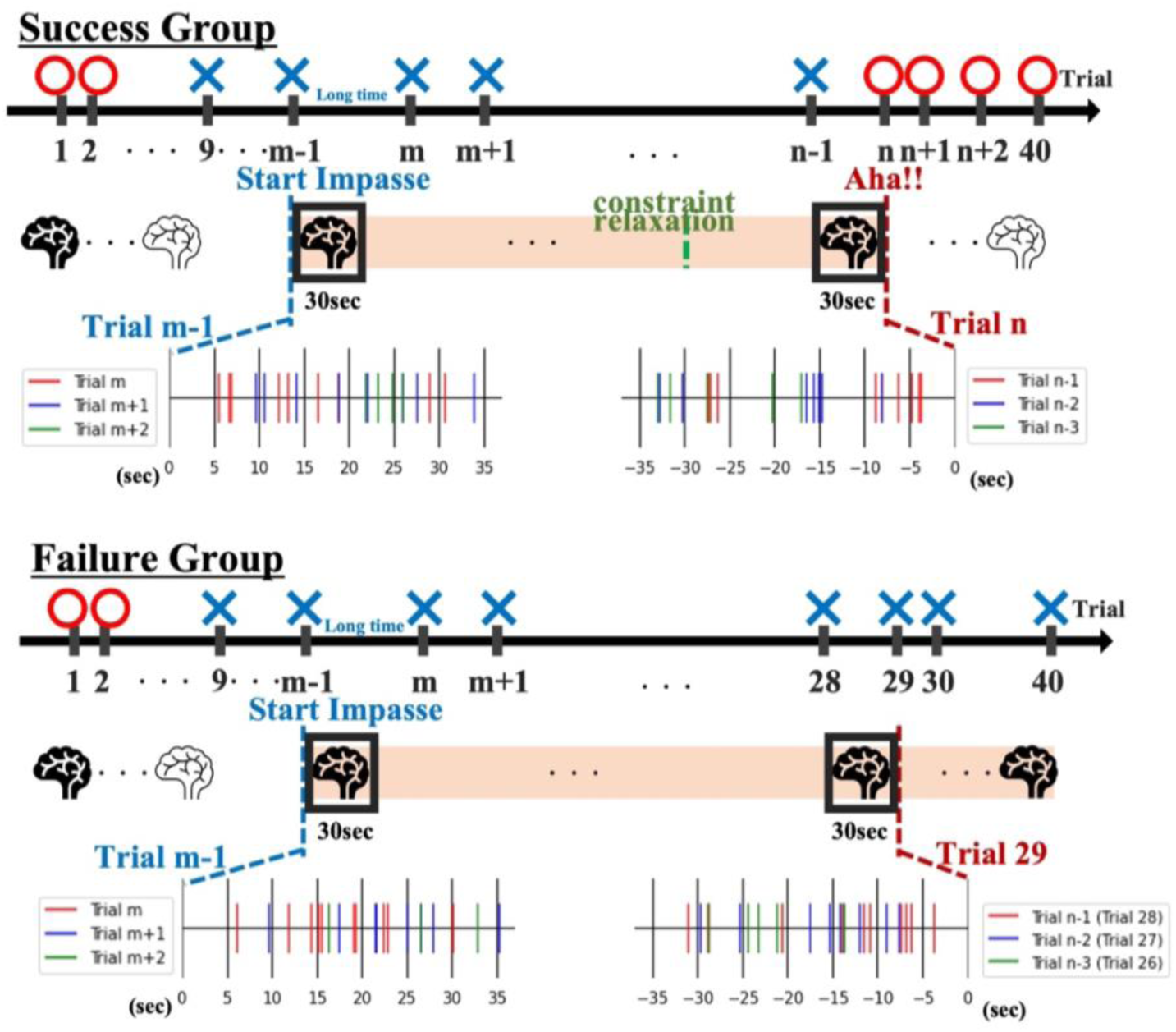
Time employed for ‘start impasse’ or ‘before aha!’ and the number of trials included in each 30 seconds. For ‘start impasse,’ 30 seconds was adopted from the start of the trial (trial m-1) before the longer considered trial (trial m). The first 30 seconds before the first trial of three consecutive correct answers was used for ‘before aha!’

### 3.2 Estimate of Constraint Relaxation by Eye Movements

We checked whether constraint relaxation occurs in the success group by increasing the ratio of vertical eye movements before the ‘aha!’ moment. The success group was averaged for the vertical ratio with trial n of each participant’s ‘aha!’ trial. The failure group was averaged over the vertical ratio during the impasse phase. For each of the n-5∼n+2 trials of the success group, a one-sample t-test was performed with the mean of the vertical ratio of the failure group. The test results revealed that the difference was significant (p<0.05, corrected by Bonferroni) for all trials from n-5∼n+2. These results indicate that the success group had a higher ratio of vertical eye movements than the failure group, even before reaching the ‘aha!’ moment. In other words, we found that the success group relaxed the constraint that the rule had to be a horizontal rule even before the trial where an ‘aha!’ moment was experienced (Figure 6).

**Figure 6.**
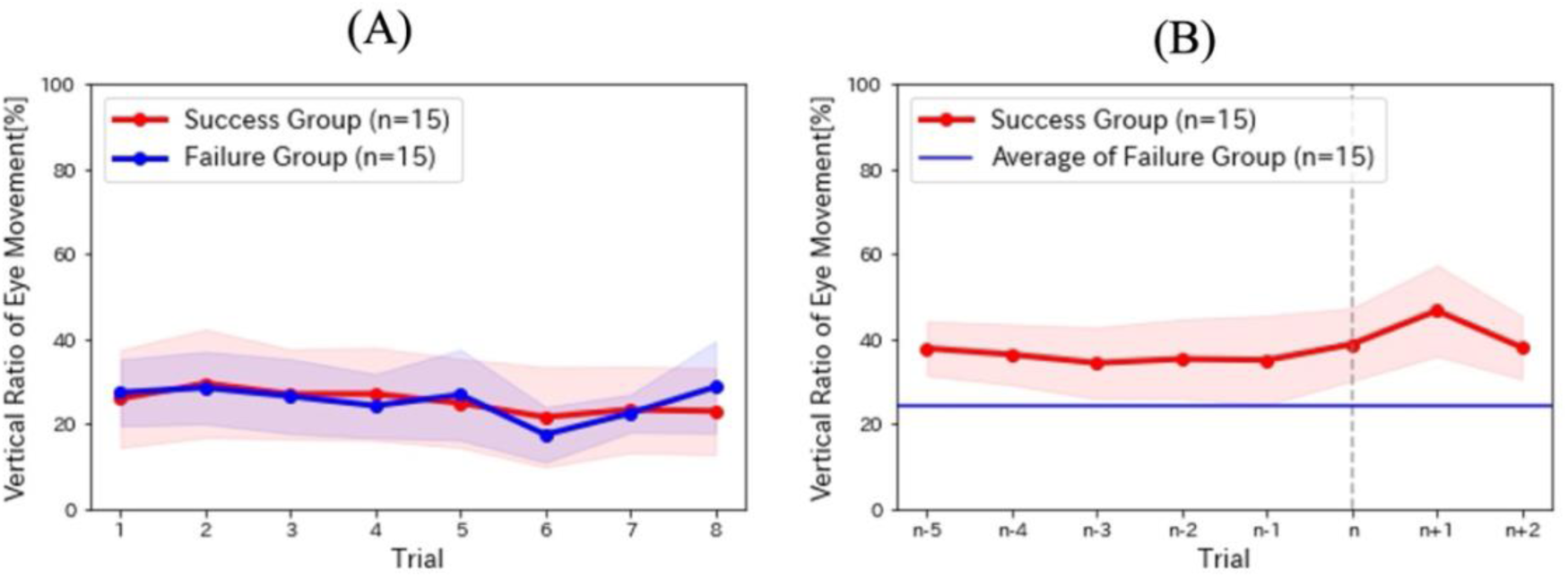
Mean vertical rate of eye movement in the success and failure groups. Red areas show 95% confidence intervals. (A) Trial 1-8 in the solidification phase. (B) Trial n-5 ∼ n+2 with ‘aha!’ as trial n.

### 3.3 Constraint Relaxation from fNIRS

In the success group, the means of 30 seconds each for ‘start impasse’ and ‘before aha!’ were compared to the 30-second mean of ‘control.’ ‘Start impasse’ activated ch41 (BA20, right inferior temporal gyrus, *t*=2.651, *p*=0.020) compared to ‘control.’ ‘Before aha!’ activated ch27 (BA46, right DLPFC, *t*=2.540, *p*=0.024) and ch41 (BA20, right inferior temporal gyrus, *t*=2.500, *p*=0.027) and deactivated ch33 (BA46, DLPFC, *t*=-2.371, *p*=0.034) compared to ‘control.’ ‘Start impasse’ and ‘before aha!’ for the successful group were compared. ‘Before aha!’ showed an activation of ch27 (BA46, right DLPFC, *t*=2.227, *p*=0.044) and ch38 (BA22, right superior temporal gyrus, *t*=2.368, *p*=0.034) compared to ‘start impasse’ (Figure 7, Appendix Table A2-4).

**Figure 7.**
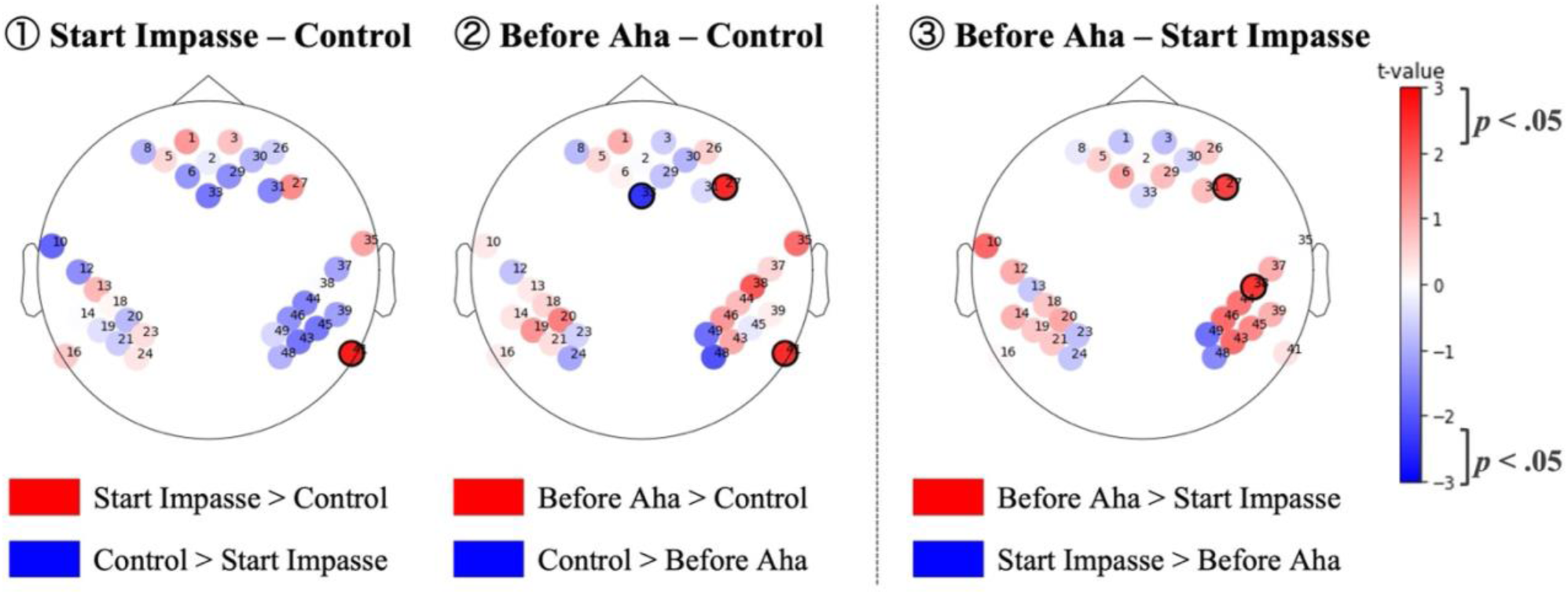
Heatmap comparing brain regions of successful groups. Black circles represent areas where there were significant differences.

In the failure group, the means of 30 seconds each of ‘start impasse’ and trial 29 were compared to the 30-second mean of ‘control.’ ‘Start impasse’ and ‘control’ were compared, but there was no significant difference. Trial 29 activated ch5 (BA10, Left Frontopolar area, *t*=2.201, *p*=0.046) compared to ‘control.’ ‘Start impasse’ and trial 29 of the failure group were compared. Trial 29 showed an activation of ch49 (BA39, right angular gyrus, right temporoparietal junction, *t*=2.694, *p*=0.018) compared to ‘start impasse’ (Figure 8, Appendix Table A5-7).

**Figure 8.**
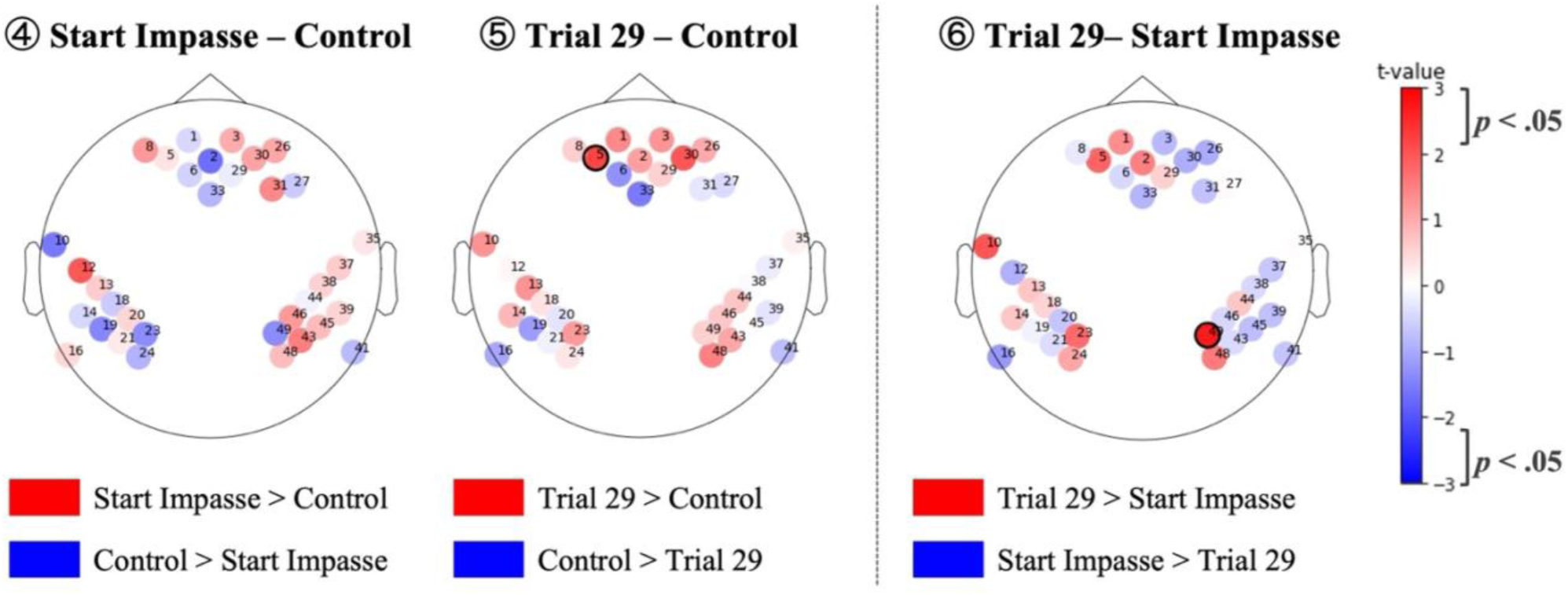
Heat map comparing brain regions in the failure group. Black circles represent areas where there were significant differences.

## 4 Discussion and Conclusion

In this study, we revealed brain activity during the constraint relaxation process in insight problem-solving. The results showed that the right DLPFC, right STG, and right ITG were involved in constraint relaxation.

Constraint relaxation was observed in the success group by measuring eye movements during the slot machine task. The success group relaxed the ‘horizontal rule’ constraint and examined the vertical rule even before the trial wherein the ‘‘aha!’ moment occurred. The same result, showing that the vertical rule had been considered before the ‘aha!’ moment, was found in Terai *et al*. (2005). A previous study on this task combined with verbal reports and eye movements revealed that relaxation was unconscious because the solver can find the correct answer as soon as the constraints are relaxed and a conscious search for the vertical rule is made (Terai, Miwa and Koga, 2005). We consider that constraint relaxation occurred unconsciously before the ‘aha!’ moment.

The participants who were able to solve insight problems activated the right STG (ch38, BA22) and right DLPFC when the constraints were relaxed. The right STG has long been reported to be involved in the ‘aha!’ phenomenon (Jung-Beeman *et al*., 2004). In our study, it was also activated because of the involvement of the ‘aha!’ experience. However, activation of the right STG has also been reported in RAT (Knyazev *et al*., 2021). We, therefore, consider the right STG to be strongly involved with constraint relaxation in the insight problem-solving process.

The DLPFC is strongly involved in cognition and memory (Barbey, Koenigs and Grafman, 2013), and the right DLPFC is related to visual working memory (Dye *et al*., 1998). Constraint relaxation requires appropriate feedback processing for each trial. On one hand, the success group may have used visual working memory through the right DLPFC to relax constraints. On the other hand, the right angular gyrus was activated in the failure group. However, as the angular gyrus (BA39, ch49) is associated with attention to salient features (Arsalidou and Taylor, 2011; Seghier, 2013), participants in the failure group may have continued to pay attention to the horizontal rule without relaxing the constraint from the horizontal to the vertical direction.

The right ITG (BA20, ch41) was activated in participants who were able to finally solve the task when they felt impasse. However, those who could not solve the task eventually showed brain activity that did not change from the previous fixation phase. The right ITG is involved in information processing for attentional recognition of visual features (Denys *et al*., 2004), and appropriate attention sharing was necessary for successful task completion.

In sum, it is possible that the fNIRS measurement of brain activity is an important indicator of constraint relaxation in the insight problem-solving process. Previously, constraint relaxation was measured by eye movements and trial-and-error behavior (Günther Knoblich, Stellan Ohlsson, 2001; Huang, Liu and Chen, 2019). Using the same slot machine task in this study, Terai *et al*. (2005) measured which hypotheses participants had in mind based on the direction of eye movements. Huang *et al*. (2019) observed constraint relaxation in RAT with reference to the order in which the participants viewed the problems to be presented on the screen. In the T-puzzle problem (make a ‘T’ out of the given blocks), constraint relaxation was determined by how the corners of the parts were used. In this study, we found that constraint relaxation might activate the right DLPFC and right STG. Brain activity, without eye movements or trial-and-error behavior, leads to the measurement of unconscious constraint relaxation, which is difficult to observe.

In conclusion, this study measured brain activity using fNIRS with the purpose of revealing the constraint relaxation process in insight problem-solving. The results suggest the involvement of the right DLPFC, right STG, and right ITG. While the method to measure brain changes caused by constraint relaxation using fNIRS can help advance related studies, including ‘aha!’ research and brain creativity research, the current study still presents limitations. First, it did not find a causal relationship between constraint relaxation and brain activity. Therefore, in the future, it is necessary to clarify the causal relationship by stimulating the brain. Salvi *et al*. (2020). showed that tDCS stimulation of the anterior temporal lobe (ATL) significantly increased RAT task performance. Aihara *et al*. (2017) stimulated the ATL with tDCS, but participants’ performance was not significantly different in either the matchstick or anagram tasks. We consider that ATL is involved in semantic comprehension (Jackson, 2021), and that stimulation of ATL promoted chunk decomposition regarding meaning, but did not affect constraint relaxation. Therefore, it is necessary to clarify the causal relationship between constraint relaxation and brain activity by stimulating the brain regions identified in this study.

Brain data is noisy and should be additionally averaged. However, for insight problem-solving, additive averaging is difficult because once ‘aha!’ occurs, the solver does not feel impasse. The same problem can be used as a case for 9-point and T-puzzle problems. We averaged the time period to be analyzed because constraint relaxation is a possible time range to some degree. In addition, we did not correct for the t-test because the data were not adequately denoised without averaging. Previous studies that measured brain activity during insight problem-solving were the anagram task and matchstick task, in which multiple problems can be prepared. However, it is not yet clear whether constraint relaxation is observed in these tasks using physiological indicators. In future, the relationship between the anagram and matchstick tasks and constraint relaxation should be clarified.

## 5 Data availability

The datasets generated and analyzed during the current study are available from the corresponding author on reasonable request.

## 6 Ethics Statement

The study involving human participants was reviewed and approved by the Ethics Review Committee of Waseda University (2020-177). The participants provided their written informed consent to participate in this study. Written informed consent was obtained from the individual(s) for the publication of any potentially identifiable images or data included in this article.

## 7 Author Contributions

R.Ohkuma and R.Osu designed the experiments and prepared the manuscript. R.Ohkuma collected and analyzed the data. Y.Kurihara, T.Takahashi and R.Osu provided important suggestions on the data analysis and the manuscript. R.Ohkuma., T.Takahashi. and R.Osu. wrote the paper.

## 9 Acknowledgments

We would like to thank Editage (www.editage.com) for English language editing.

