## Appendices for "Brain Activity During Constraint Relaxation in the Insight Problem-Solving Process: An fNIRS Study"

##### Table A1 Numbers displayed in the slot machine task in the main task.

| Trial | Slot 1 | Slot 2 | Slot 3 | Dummy Rule |
|---|---|---|---|---|
| 1 | 3 | 4 | 7 | ○ |
| 2 | 4 | 6 | 0 | ○ |
| 3 | 9 | 4 | 3 | ○ |
| 4 | 4 | 2 | 6 | ○ |
| 5 | 3 | 6 | 9 | ○ |
| 6 | 2 | 0 | 2 | ○ |
| 7 | 2 | 3 | 5 | ○ |
| 8 | 5 | 3 | 8 | ○ |
| 9 | 0 | 2 | 1 | |
| 10 | 2 | 2 | 4 | ○ |
| 11 | 6 | 3 | 7 | |
| 12 | 2 | 8 | 0 | ○ |
| 13 | 3 | 0 | 3 | ○ |
| 14 | 5 | 3 | 6 | |
| 15 | 4 | 4 | 9 | |
| 16 | 4 | 8 | 2 | ○ |
| 17 | 3 | 5 | 5 | |
| 18 | 3 | 1 | 8 | |
| 19 | 0 | 1 | 1 | ○ |
| 20 | 6 | 2 | 4 | |
| 21 | 5 | 2 | 7 | ○ |
| 22 | 0 | 0 | 0 | ○ |
| 23 | 2 | 7 | 3 | |
| 24 | 7 | 3 | 6 | |
| 25 | 3 | 5 | 9 | |
| 26 | 5 | 6 | 2 | |
| 27 | 3 | 2 | 5 | ○ |
| 28 | 3 | 2 | 8 | |
| 29 | 4 | 6 | 1 | |
| 30 | 2 | 6 | 4 | |
| 31 | 3 | 3 | 7 | |
| 32 | 6 | 3 | 0 | |
| 33 | 8 | 1 | 3 | |
| 34 | 5 | 5 | 6 | |
| 35 | 4 | 7 | 9 | |
| 36 | 3 | 6 | 2 | |
| 37 | 7 | 5 | 5 | |
| 38 | 5 | 8 | 8 | |
| 39 | 3 | 7 | 1 | |
| 40 | 4 | 4 | 4 | |

### Table A2 t-test results. ① 'Start Impasse' – 'Control' (Success Groups)

| Brodman | Region name | | channel | Control | | Before Aha | | | |
|---|---|---|---|---|---|---|---|---|---|
| | | | | acerage | *SD* | average | *SD* | *t* | *p* |
| BA10 | Frontopolar area | | ch2 | 0.000 | 0.005 | -0.001 | 0.005 | -0.229 | 0.823 |
| | | L | ch5 | 0.000 | 0.003 | 0.000 | 0.004 | 0.447 | 0.663 |
| | | | ch6 | 0.002 | 0.006 | -0.001 | 0.003 | -1.239 | 0.237 |
| | | R | ch29 | 0.002 | 0.004 | -0.001 | 0.003 | -1.313 | 0.212 |
| | | | ch30 | 0.001 | 0.004 | -0.001 | 0.003 | -0.880 | 0.395 |
| BA11 | Orbitofrontal area | L | ch1 | -0.017 | 0.055 | -0.001 | 0.017 | 1.199 | 0.252 |
| | | | ch8 | 0.025 | 0.090 | 0.004 | 0.013 | -0.895 | 0.387 |
| | | R | ch3 | 0.000 | 0.005 | 0.001 | 0.008 | 0.707 | 0.492 |
| | | | ch26 | -0.001 | 0.003 | -0.001 | 0.005 | -0.584 | 0.569 |
| BA9, 46 | Dorsolateral prefrontal cortex | | ch33 | 0.003 | 0.006 | -0.001 | 0.004 | -1.602 | 0.133 |
| | | R | ch27 | 0.000 | 0.003 | 0.001 | 0.002 | 1.373 | 0.193 |
| | | | ch31 | 0.002 | 0.004 | 0.000 | 0.003 | -1.415 | 0.181 |
| BA20 | Inferior Temporal gyrus | L | ch14 | 0.000 | 0.007 | 0.000 | 0.006 | -0.036 | 0.972 |
| | | | ch16 | 0.001 | 0.011 | 0.004 | 0.014 | 0.623 | 0.544 |
| | | R | ch39 | 0.001 | 0.002 | -0.001 | 0.005 | -1.115 | 0.285 |
| | | | ch41* | -0.004 | 0.006 | 0.001 | 0.005 | 2.651 | 0.020 |
| BA21 | Middle Temporal gyrus | L | ch10 | 0.003 | 0.007 | -0.002 | 0.005 | -1.812 | 0.093 |
| | | | ch12 | 0.001 | 0.005 | -0.002 | 0.008 | -1.333 | 0.205 |
| | | R | ch35 | -0.003 | 0.008 | 0.000 | 0.005 | 1.099 | 0.292 |
| | | | ch37 | 0.000 | 0.002 | -0.001 | 0.003 | -1.072 | 0.303 |
| BA22 | Superior Temporal gyrus | L | ch13 | -0.001 | 0.010 | 0.001 | 0.012 | 0.829 | 0.422 |
| | | | ch18 | 0.000 | 0.008 | 0.000 | 0.003 | 0.096 | 0.925 |
| | | | ch19 | 0.001 | 0.009 | 0.000 | 0.004 | -0.371 | 0.717 |
| | | R | ch38 | 0.000 | 0.003 | 0.000 | 0.003 | -0.037 | 0.971 |
| | | | ch44 | 0.002 | 0.005 | -0.001 | 0.003 | -1.447 | 0.172 |
| | | | ch45 | 0.002 | 0.006 | -0.001 | 0.003 | -1.600 | 0.134 |
| BA39 | Angular gyrus / | L | ch20 | 0.002 | 0.009 | -0.001 | 0.003 | -0.814 | 0.430 |
| | Temporo-parietal junction | | ch21 | 0.001 | 0.008 | 0.000 | 0.003 | -0.609 | 0.553 |
| | | | ch23 | -0.001 | 0.004 | -0.001 | 0.004 | 0.391 | 0.702 |
| | | | ch24 | -0.001 | 0.004 | -0.001 | 0.003 | 0.291 | 0.776 |
| | | R | ch43 | 0.001 | 0.006 | -0.002 | 0.004 | -1.601 | 0.133 |
| | | | ch46 | 0.001 | 0.007 | -0.002 | 0.004 | -1.277 | 0.224 |
| | | | ch48 | 0.000 | 0.002 | -0.001 | 0.003 | -0.900 | 0.385 |
| | | | ch49 | 0.002 | 0.009 | 0.001 | 0.003 | -0.457 | 0.655 |

(*$p < .05$)

#### Table A3 t-test results. ② 'Before Aha!' – 'Control' (Success Groups)

| Brodman | Region name | | channel | Control acerage | Control *SD* | Before Aha average | Before Aha *SD* | *t* | *p* |
|---|---|---|---|---|---|---|---|---|---|
| BA10 | Frontopolar area | | ch2 | 0.000 | 0.005 | 0.000 | 0.004 | -0.064 | 0.950 |
| | | L | ch5 | 0.000 | 0.004 | 0.002 | 0.015 | 0.460 | 0.653 |
| | | | ch6 | 0.002 | 0.007 | 0.003 | 0.011 | 0.141 | 0.890 |
| | | R | ch29 | 0.002 | 0.005 | 0.001 | 0.006 | -0.714 | 0.488 |
| | | | ch30 | 0.001 | 0.004 | -0.002 | 0.007 | -0.873 | 0.399 |
| BA11 | Orbitofrontal area | L | ch1 | -0.021 | 0.070 | 0.008 | 0.039 | 0.961 | 0.354 |
| | | | ch8 | 0.026 | 0.096 | 0.002 | 0.019 | -0.815 | 0.430 |
| | | R | ch3 | -0.001 | 0.008 | -0.003 | 0.009 | -0.604 | 0.556 |
| | | | ch26 | 0.000 | 0.003 | 0.000 | 0.005 | 0.531 | 0.604 |
| BA9, 46 | Dorsolateral prefrontal cortex | | ch33* | 0.004 | 0.006 | -0.001 | 0.007 | -2.371 | 0.034 |
| | | R | ch27* | 0.000 | 0.003 | 0.003 | 0.004 | 2.540 | 0.025 |
| | | | ch31 | 0.003 | 0.005 | 0.002 | 0.008 | -0.445 | 0.664 |
| BA20 | Inferior Temporal gyrus | L | ch14 | -0.001 | 0.008 | 0.000 | 0.006 | 0.320 | 0.754 |
| | | | ch16 | 0.002 | 0.016 | 0.003 | 0.017 | 0.228 | 0.823 |
| | | R | ch39 | 0.001 | 0.003 | 0.001 | 0.005 | 0.195 | 0.848 |
| | | | ch41* | -0.004 | 0.007 | 0.000 | 0.006 | 2.496 | 0.027 |
| BA21 | Middle Temporal gyrus | L | ch10 | 0.003 | 0.007 | 0.003 | 0.008 | 0.260 | 0.799 |
| | | | ch12 | 0.002 | 0.005 | 0.000 | 0.006 | -0.703 | 0.494 |
| | | R | ch35 | -0.005 | 0.012 | 0.000 | 0.005 | 1.725 | 0.108 |
| | | | ch37 | 0.000 | 0.002 | 0.001 | 0.005 | 0.474 | 0.643 |
| BA22 | Superior Temporal gyrus | L | ch13 | 0.000 | 0.010 | 0.000 | 0.004 | 0.252 | 0.805 |
| | | | ch18 | 0.000 | 0.009 | 0.001 | 0.008 | 0.496 | 0.628 |
| | | | ch19 | 0.002 | 0.010 | 0.003 | 0.013 | 1.263 | 0.229 |
| | | R | ch38 | 0.001 | 0.003 | 0.003 | 0.004 | 1.987 | 0.068 |
| | | | ch44 | 0.001 | 0.006 | 0.004 | 0.011 | 0.797 | 0.440 |
| | | | ch45 | 0.002 | 0.007 | 0.002 | 0.007 | -0.261 | 0.798 |
| BA39 | Angular gyrus / | L | ch20 | 0.002 | 0.011 | 0.004 | 0.013 | 1.507 | 0.156 |
| | Temporo-parietal junction | | ch21 | 0.002 | 0.009 | 0.002 | 0.011 | 0.378 | 0.711 |
| | | | ch23 | -0.001 | 0.004 | -0.002 | 0.003 | -0.521 | 0.611 |
| | | | ch24 | -0.001 | 0.004 | -0.002 | 0.004 | -1.033 | 0.321 |
| | | R | ch43 | 0.001 | 0.007 | 0.007 | 0.017 | 1.022 | 0.325 |
| | | | ch46 | 0.000 | 0.008 | 0.006 | 0.015 | 1.207 | 0.249 |
| | | | ch48 | 0.000 | 0.002 | -0.004 | 0.008 | -2.062 | 0.060 |
| | | | ch49 | 0.002 | 0.010 | -0.005 | 0.010 | -1.708 | 0.111 |

*(*p < .05)*

#### Table A4 t-test results. ③ 'Before Aha!' – 'Start Impasse' (Success Groups)

| Brodman | Region name | | channel | Control acerage | Control *SD* | Before Aha average | Before Aha *SD* | *t* | *p* |
|---|---|---|---|---|---|---|---|---|---|
| BA10 | Frontopolar area | | ch2 | 0.000 | 0.006 | 0.000 | 0.004 | 0.052 | 0.960 |
| | | L | ch5 | -0.001 | 0.004 | 0.002 | 0.015 | 0.551 | 0.591 |
| | | | ch6 | -0.001 | 0.005 | 0.003 | 0.011 | 1.046 | 0.314 |
| | | R | ch29 | -0.001 | 0.006 | 0.001 | 0.006 | 0.782 | 0.448 |
| | | | ch30 | -0.001 | 0.006 | -0.002 | 0.007 | -0.479 | 0.640 |
| BA11 | Orbitofrontal area | L | ch1 | 0.019 | 0.054 | 0.008 | 0.039 | -0.694 | 0.500 |
| | | | ch8 | 0.004 | 0.020 | 0.002 | 0.019 | -0.279 | 0.785 |
| | | R | ch3 | -0.001 | 0.009 | -0.003 | 0.009 | -0.787 | 0.445 |
| | | | ch26 | -0.001 | 0.005 | 0.000 | 0.005 | 0.632 | 0.538 |
| BA9, 46 | Dorsolateral prefrontal cortex | | ch33 | 0.000 | 0.004 | -0.001 | 0.007 | -0.443 | 0.665 |
| | | R | ch27* | 0.000 | 0.003 | 0.003 | 0.004 | 2.227 | 0.044 |
| | | | ch31 | 0.000 | 0.004 | 0.002 | 0.008 | 0.771 | 0.454 |
| BA20 | Inferior Temporal gyrus | L | ch14 | -0.004 | 0.013 | 0.000 | 0.006 | 0.928 | 0.370 |
| | | | ch16 | 0.003 | 0.015 | 0.003 | 0.017 | 0.087 | 0.932 |
| | | R | ch39 | -0.001 | 0.005 | 0.001 | 0.005 | 0.917 | 0.376 |
| | | | ch41 | 0.000 | 0.005 | 0.000 | 0.006 | 0.346 | 0.735 |
| BA21 | Middle Temporal gyrus | L | ch10 | -0.002 | 0.006 | 0.003 | 0.008 | 1.789 | 0.097 |
| | | | ch12 | -0.003 | 0.009 | 0.000 | 0.006 | 0.975 | 0.347 |
| | | R | ch35 | 0.000 | 0.006 | 0.000 | 0.005 | -0.002 | 0.998 |
| | | | ch37 | -0.001 | 0.004 | 0.001 | 0.005 | 0.948 | 0.361 |
| BA22 | Superior Temporal gyrus | L | ch13 | 0.002 | 0.009 | 0.000 | 0.004 | -0.670 | 0.514 |
| | | | ch18 | -0.001 | 0.003 | 0.001 | 0.008 | 0.674 | 0.512 |
| | | | ch19 | 0.000 | 0.005 | 0.003 | 0.013 | 0.645 | 0.530 |
| | | R | ch38* | 0.000 | 0.003 | 0.003 | 0.004 | 2.368 | 0.034 |
| | | | ch44 | -0.002 | 0.004 | 0.004 | 0.011 | 1.637 | 0.126 |
| | | | ch45 | -0.002 | 0.006 | 0.002 | 0.007 | 1.394 | 0.187 |
| BA39 | Angular gyrus / | L | ch20 | -0.001 | 0.003 | 0.004 | 0.013 | 1.022 | 0.325 |
| | Temporo-parietal junction | | ch21 | 0.000 | 0.003 | 0.002 | 0.011 | 0.649 | 0.528 |
| | | | ch23 | 0.000 | 0.004 | -0.002 | 0.003 | -0.736 | 0.475 |
| | | | ch24 | -0.001 | 0.003 | -0.002 | 0.004 | -0.692 | 0.501 |
| | | R | ch43 | -0.003 | 0.005 | 0.007 | 0.017 | 1.737 | 0.106 |
| | | | ch46 | -0.003 | 0.005 | 0.006 | 0.015 | 1.726 | 0.108 |
| | | | ch48 | 0.000 | 0.004 | -0.004 | 0.008 | -1.385 | 0.189 |
| | | | ch49 | 0.001 | 0.004 | -0.005 | 0.010 | -1.686 | 0.116 |

(*$p < .05$)

#### Table A5 t-test results. ④ 'Start Impasse' – 'Control' (Failure Groups)

| Brodman | Region name | | channel | Control | | Before Aha | | | |
|---|---|---|---|---|---|---|---|---|---|
| | | | | acerage | *SD* | average | *SD* | *t* | *p* |
| BA10 | Frontopolar area | | ch2 | 0.001 | 0.003 | -0.001 | 0.005 | -1.692 | 0.114 |
| | | L | ch5 | -0.001 | 0.002 | 0.000 | 0.002 | 0.306 | 0.765 |
| | | | ch6 | 0.000 | 0.002 | -0.001 | 0.003 | -0.527 | 0.607 |
| | | R | ch29 | 0.000 | 0.001 | 0.000 | 0.001 | -0.222 | 0.828 |
| | | | ch30 | -0.003 | 0.008 | 0.018 | 0.062 | 1.062 | 0.308 |
| BA11 | Orbitofrontal area | L | ch1 | 0.000 | 0.002 | -0.001 | 0.006 | -0.451 | 0.660 |
| | | | ch8 | 0.000 | 0.002 | 0.001 | 0.005 | 1.210 | 0.248 |
| | | R | ch3 | -0.003 | 0.010 | 0.010 | 0.038 | 0.983 | 0.344 |
| | | | ch26 | -0.003 | 0.009 | 0.028 | 0.102 | 1.048 | 0.314 |
| BA9, 46 | Dorsolateral prefrontal cortex | | ch33 | 0.000 | 0.003 | 0.000 | 0.003 | -0.870 | 0.400 |
| | | R | ch27 | 0.000 | 0.006 | -0.001 | 0.004 | -0.619 | 0.547 |
| | | | ch31 | 0.000 | 0.001 | 0.001 | 0.001 | 1.403 | 0.184 |
| BA20 | Inferior Temporal gyrus | L | ch14 | 0.002 | 0.003 | 0.000 | 0.008 | -0.463 | 0.651 |
| | | | ch16 | 0.002 | 0.012 | 0.004 | 0.017 | 0.459 | 0.654 |
| | | R | ch39 | 0.000 | 0.002 | 0.000 | 0.002 | 0.500 | 0.626 |
| | | | ch41 | 0.000 | 0.009 | -0.002 | 0.006 | -0.799 | 0.439 |
| BA21 | Middle Temporal gyrus | L | ch10 | 0.001 | 0.005 | -0.002 | 0.005 | -1.591 | 0.136 |
| | | | ch12 | 0.000 | 0.002 | 0.001 | 0.003 | 1.988 | 0.068 |
| | | R | ch35 | 0.001 | 0.004 | 0.001 | 0.006 | 0.300 | 0.769 |
| | | | ch37 | 0.000 | 0.003 | 0.000 | 0.003 | 0.609 | 0.553 |
| BA22 | Superior Temporal gyrus | L | ch13 | 0.001 | 0.002 | 0.002 | 0.005 | 0.603 | 0.557 |
| | | | ch18 | 0.000 | 0.003 | 0.000 | 0.004 | -0.617 | 0.548 |
| | | | ch19 | 0.001 | 0.004 | 0.000 | 0.004 | -1.427 | 0.177 |
| | | R | ch38 | 0.000 | 0.002 | 0.001 | 0.003 | 0.530 | 0.605 |
| | | | ch44 | 0.000 | 0.002 | 0.000 | 0.003 | -0.141 | 0.890 |
| | | | ch45 | 0.000 | 0.002 | 0.001 | 0.003 | 0.808 | 0.434 |
| BA39 | Angular gyrus / | L | ch20 | -0.001 | 0.006 | 0.001 | 0.006 | 0.474 | 0.644 |
| | Temporo-parietal junction | | ch21 | 0.000 | 0.006 | 0.000 | 0.002 | 0.245 | 0.811 |
| | | | ch23 | 0.000 | 0.003 | -0.002 | 0.005 | -1.340 | 0.203 |
| | | | ch24 | 0.001 | 0.004 | -0.001 | 0.006 | -0.904 | 0.383 |
| | | R | ch43 | -0.002 | 0.005 | 0.002 | 0.005 | 1.506 | 0.156 |
| | | | ch46 | -0.001 | 0.003 | 0.001 | 0.004 | 1.194 | 0.254 |
| | | | ch48 | 0.000 | 0.003 | 0.001 | 0.004 | 0.778 | 0.450 |
| | | | ch49 | 0.001 | 0.004 | -0.004 | 0.010 | -1.342 | 0.203 |

*(*p < .05)*

#### Table A6 t-test results. ⑤ Trial 29 – 'Control' (Failure Groups)

| Brodman | Region name | | channel | Control acerage | Control *SD* | Before Aha average | Before Aha *SD* | *t* | *p* |
|---|---|---|---|---|---|---|---|---|---|
| BA10 | Frontopolar area | | ch2 | 0.001 | 0.003 | 0.004 | 0.009 | 1.114 | 0.285 |
| | | L | ch5* | -0.001 | 0.002 | 0.002 | 0.004 | 2.201 | 0.046 |
| | | | ch6 | 0.000 | 0.002 | -0.001 | 0.002 | -1.269 | 0.227 |
| | | R | ch29 | 0.000 | 0.001 | 0.000 | 0.002 | 0.553 | 0.590 |
| | | | ch30 | -0.003 | 0.008 | -0.001 | 0.009 | 1.975 | 0.070 |
| BA11 | Orbitofrontal area | L | ch1 | 0.000 | 0.002 | 0.003 | 0.006 | 1.385 | 0.191 |
| | | | ch8 | 0.000 | 0.002 | 0.001 | 0.008 | 0.586 | 0.568 |
| | | R | ch3 | -0.003 | 0.010 | 0.003 | 0.006 | 1.259 | 0.230 |
| | | | ch26 | -0.003 | 0.009 | -0.001 | 0.007 | 0.990 | 0.340 |
| BA9, 46 | Dorsolateral prefrontal cortex | | ch33 | 0.000 | 0.003 | -0.001 | 0.003 | -1.562 | 0.142 |
| | | R | ch27 | 0.000 | 0.006 | -0.001 | 0.004 | -0.444 | 0.665 |
| | | | ch31 | 0.000 | 0.001 | -0.001 | 0.005 | -0.301 | 0.768 |
| BA20 | Inferior Temporal gyrus | L | ch14 | 0.002 | 0.003 | 0.003 | 0.007 | 0.840 | 0.416 |
| | | | ch16 | 0.002 | 0.012 | -0.006 | 0.015 | -1.046 | 0.315 |
| | | R | ch39 | 0.000 | 0.002 | 0.000 | 0.003 | -0.325 | 0.750 |
| | | | ch41 | 0.000 | 0.009 | -0.007 | 0.028 | -0.753 | 0.465 |
| BA21 | Middle Temporal gyrus | L | ch10 | 0.001 | 0.005 | 0.004 | 0.011 | 1.279 | 0.223 |
| | | | ch12 | 0.000 | 0.002 | 0.000 | 0.004 | 0.108 | 0.915 |
| | | R | ch35 | 0.001 | 0.004 | 0.001 | 0.014 | 0.173 | 0.865 |
| | | | ch37 | 0.000 | 0.003 | -0.001 | 0.004 | -0.221 | 0.828 |
| BA22 | Superior Temporal gyrus | L | ch13 | 0.001 | 0.002 | 0.003 | 0.008 | 1.269 | 0.227 |
| | | | ch18 | 0.000 | 0.003 | 0.001 | 0.005 | 0.329 | 0.747 |
| | | | ch19 | 0.001 | 0.004 | 0.000 | 0.004 | -1.153 | 0.270 |
| | | R | ch38 | 0.000 | 0.002 | 0.000 | 0.005 | -0.069 | 0.946 |
| | | | ch44 | 0.000 | 0.002 | 0.001 | 0.004 | 0.673 | 0.513 |
| | | | ch45 | 0.000 | 0.002 | 0.000 | 0.003 | 0.046 | 0.964 |
| BA39 | Angular gyrus / | L | ch20 | -0.001 | 0.006 | -0.001 | 0.005 | -0.323 | 0.752 |
| | Temporo-parietal junction | | ch21 | 0.000 | 0.006 | -0.001 | 0.006 | -0.167 | 0.870 |
| | | | ch23 | 0.000 | 0.003 | 0.002 | 0.004 | 1.178 | 0.260 |
| | | | ch24 | 0.001 | 0.004 | 0.001 | 0.003 | 0.302 | 0.768 |
| | | R | ch43 | -0.002 | 0.005 | 0.001 | 0.006 | 1.041 | 0.317 |
| | | | ch46 | -0.001 | 0.003 | 0.000 | 0.005 | 0.651 | 0.526 |
| | | | ch48 | 0.000 | 0.003 | 0.004 | 0.006 | 1.496 | 0.159 |
| | | | ch49 | 0.001 | 0.004 | 0.003 | 0.011 | 0.543 | 0.596 |

*(*p < .05)*

#### Table A7 t-test results. ⑥ Trial 29 – 'Start Impasse' (Failure Groups)

| Brodman | Region name | | channel | Control | | Before Aha | | | |
|---|---|---|---|---|---|---|---|---|---|
| | | | | acerage | *SD* | average | *SD* | *t* | *p* |
| BA10 | Frontopolar area | | ch2 | -0.001 | 0.005 | 0.004 | 0.009 | 1.492 | 0.160 |
| | | L | ch5 | 0.000 | 0.002 | 0.002 | 0.004 | 1.812 | 0.093 |
| | | | ch6 | -0.001 | 0.003 | -0.001 | 0.002 | -0.458 | 0.654 |
| | | R | ch29 | 0.000 | 0.001 | 0.000 | 0.002 | 0.557 | 0.587 |
| | | | ch30 | 0.018 | 0.062 | -0.001 | 0.009 | -0.958 | 0.356 |
| BA11 | Orbitofrontal area | L | ch1 | -0.001 | 0.006 | 0.003 | 0.006 | 1.325 | 0.210 |
| | | | ch8 | 0.001 | 0.005 | 0.001 | 0.008 | -0.281 | 0.783 |
| | | R | ch3 | 0.010 | 0.038 | 0.003 | 0.006 | -0.827 | 0.423 |
| | | | ch26 | 0.028 | 0.102 | -0.001 | 0.007 | -0.976 | 0.347 |
| BA9, 46 | Dorsolateral prefrontal cortex | | ch33 | 0.000 | 0.003 | -0.001 | 0.003 | -0.870 | 0.400 |
| | | R | ch27 | -0.001 | 0.004 | -0.001 | 0.004 | 0.052 | 0.959 |
| | | | ch31 | 0.001 | 0.001 | -0.001 | 0.005 | -0.707 | 0.492 |
| BA20 | Inferior Temporal gyrus | L | ch14 | 0.000 | 0.008 | 0.003 | 0.007 | 0.659 | 0.522 |
| | | | ch16 | 0.004 | 0.017 | -0.006 | 0.015 | -1.199 | 0.252 |
| | | R | ch39 | 0.000 | 0.002 | 0.000 | 0.003 | -0.732 | 0.477 |
| | | | ch41 | -0.002 | 0.006 | -0.007 | 0.028 | -0.702 | 0.495 |
| BA21 | Middle Temporal gyrus | L | ch10 | -0.002 | 0.005 | 0.004 | 0.011 | 2.009 | 0.066 |
| | | | ch12 | 0.001 | 0.003 | 0.000 | 0.004 | -0.934 | 0.367 |
| | | R | ch35 | 0.001 | 0.006 | 0.001 | 0.014 | 0.037 | 0.971 |
| | | | ch37 | 0.000 | 0.003 | -0.001 | 0.004 | -0.662 | 0.520 |
| BA22 | Superior Temporal gyrus | L | ch13 | 0.002 | 0.005 | 0.003 | 0.008 | 0.660 | 0.521 |
| | | | ch18 | 0.000 | 0.004 | 0.001 | 0.005 | 0.501 | 0.625 |
| | | | ch19 | 0.000 | 0.004 | 0.000 | 0.004 | -0.156 | 0.879 |
| | | R | ch38 | 0.001 | 0.003 | 0.000 | 0.005 | -0.447 | 0.662 |
| | | | ch44 | 0.000 | 0.003 | 0.001 | 0.004 | 0.675 | 0.511 |
| | | | ch45 | 0.001 | 0.003 | 0.000 | 0.003 | -0.755 | 0.464 |
| BA39 | Angular gyrus / | L | ch20 | 0.001 | 0.006 | -0.001 | 0.005 | -0.696 | 0.499 |
| | Temporo-parietal junction | | ch21 | 0.000 | 0.002 | -0.001 | 0.006 | -0.407 | 0.691 |
| | | | ch23 | -0.002 | 0.005 | 0.002 | 0.004 | 1.755 | 0.103 |
| | | | ch24 | -0.001 | 0.006 | 0.001 | 0.003 | 1.045 | 0.315 |
| | | R | ch43 | 0.002 | 0.005 | 0.001 | 0.006 | -0.431 | 0.674 |
| | | | ch46 | 0.001 | 0.004 | 0.000 | 0.005 | -0.478 | 0.641 |
| | | | ch48 | 0.001 | 0.004 | 0.004 | 0.006 | 1.563 | 0.142 |
| | | | ch49* | -0.004 | 0.010 | 0.003 | 0.011 | 2.694 | 0.018 |

*(*p < .05)*
